# Regnase-1 deficiency restrains *Klebsiella pneumoniae* infection by regulation of a Type I interferon response

**DOI:** 10.1101/2021.11.20.469389

**Authors:** Giraldina Trevejo-Nuñez, Becky Lin, Li Fan, Felix E. Y. Aggor, Partha S. Biswas, Kong Chen, Sarah L. Gaffen

## Abstract

Excessive inflammatory responses can cause collateral tissue damage or autoimmune inflammation, sometimes with severe morbidity or mortality. During host defense responses, numerous negative feedback mechanisms are established to prevent excessive unchecked inflammation. However, this restraint can sometimes come at the cost of suboptimal infection control, and we do not fully understand how this balance is maintained during different infection settings. The endoribonuclease Regnase-1 (Reg1, Zc3h12a, MCPIP1) is an RNA binding protein (RBP) that binds and degrades many target mRNA transcripts. Reg1 is a potent feedback regulator of IL-17 and LPS signal transduction, among other stimuli. Consequently, Reg1 deficiency exacerbates autoimmune inflammation in multiple mouse models, but on the other hand, reduced Reg1 improves immunity to fungal infection. To date, the role of Reg1 in bacterial immunity is poorly defined. Here, we show that mice deficient in Reg1 are more resistant to pulmonary *Klebsiella pneumoniae* (KP) infection. Unexpectedly, effects of Reg1 deficiency were not due to accelerated eradication of bacteria or increased pro-inflammatory cytokine expression. Rather, alveolar macrophages from Reg1-deficient mice showed enrichment of Type I IFN-related genes upon KP infection, accompanied by increased Ifnb1 expression. Surprisingly, the stability of Ifnb1 mRNA was not altered by Reg1-deficiency; rather, mRNA encoding its upstream regulator IRF7 appeared to be a more prominent target. Blockade of IFNR during KP infection reversed disease improvement. Thus, impaired Reg1 induces Type I IFN and enhances resistance to KP, raising the possibility that Reg1 could be a potential clinical target in acute bacterial infections.

**Graphical abstract:** 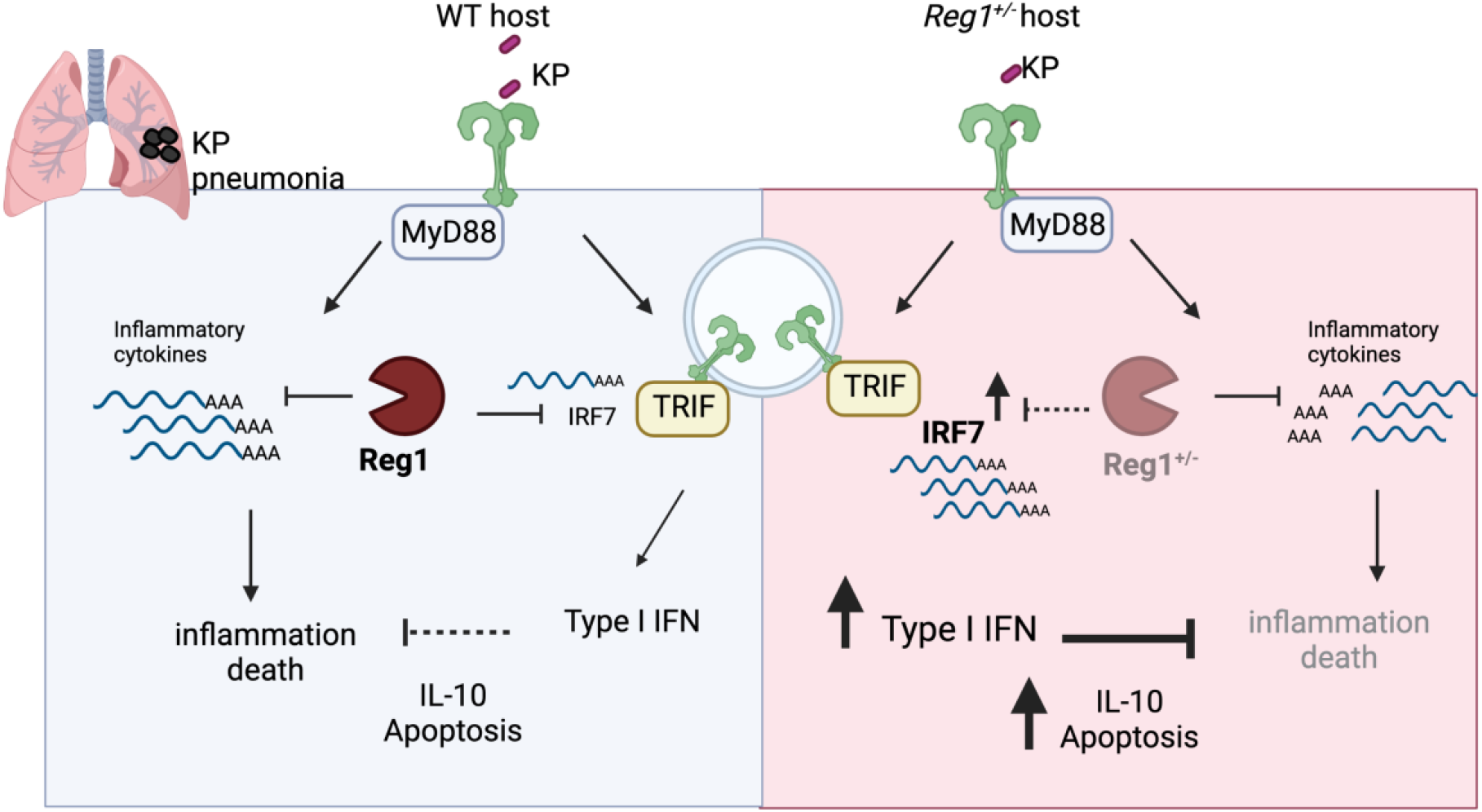

## Introduction

*Klebsiella pneumoniae* (KP) is a constituent of the normal human normal human microbiome and is detected in approximately 10% of the Human Microbiome Project samples (1, 2). KP is the third most common cause of ventilator-associated pneumonia (VAP) in the USA, and its presence correlates with prolonged duration of ventilation and ICU hospital stay (3–5). KP imposes a major infectious disease challenge due to increasing antimicrobial resistance combined with comparatively limited therapeutic options (6). Hence, an improved understanding of host-pathogen interactions is needed to elucidate, and ultimately exploit, the host mechanisms that control KP pneumonia.

The immune response to infectious pathogens is finely balanced to allow eradication of microbes yet still prevent excessive inflammatory pathology. Thus, numerous mechanisms are triggered during immune responses to counter-regulate deleterious responses and thereby minimize collateral damage to the host. The RNA-binding protein (RBP) Regnase-1 (Reg1), also known as MCP1-induced protein 1 (MCPIP1), is encoded by the *ZC3H12A* gene and has intrinsic endoribonuclease activity that keeps inflammatory responses in check (7). In this capacity, Reg1 binds and degrades many mRNA transcripts, including its own mRNA, through recognition of characteristic stem-loop secondary structures in the 3’ untranslated region (UTR) of target mRNAs (8). Reg1 is induced by many inflammatory stimuli, including IL-1, Toll-like-receptors (TLR) and IL-17 (9–12). In addition, TCR signaling leads to cleavage of Reg1, releasing T cells from Reg1-mediated suppression (13). Thus, Reg1 plays a central role in negative feedback control of pro-inflammatory cytokine gene expression through post-transcriptional control of mRNA expression.

Multiple mouse models support the vital role of Reg1 as a central negative regulator of pathogenic inflammation. Reg1 knockout mice (*Zc3h12a*^−/−^, here called *Reg1*^−/−^) have a life span only of 6-8 weeks. *Reg1*^−/−^ mice display significant constitutive inflammation, manifesting as growth retardation, splenomegaly, lymphadenopathy, heightened systemic and localized cytokine production, and ultimately multi-organ failure (9, 14). In order to avoid confounding issues with the complete Reg1 knockout mouse, we have previously taken advantage of *Reg1* heterozygous mice (*Reg1^+/−^*), which have normal life spans and fertility, normal tonic expression of Reg1, and minimal peripheral organ inflammation at baseline (11, 15). Even so, cells from *Reg1^+/−^* mice display markedly reduced Reg1 expression after LPS or IL-17 stimulation, and tissues from *Reg1^+/−^* animals (lung, kidney, central nervous system) exhibit elevated cytokine responses (11). Consequently, *Reg1^+/−^* mice show exacerbated signs of autoimmunity in multiple IL-17-driven autoimmune conditions, including experimental autoimmune encephalomyelitis (EAE), imiquimod (IMQ)-induced dermatitis and autoantibody-induced glomerulonephritis (AGN). In all these settings, IL-17R deficiency reversed the phenotype, demonstrating that Reg1 functions in these settings through restricting IL-17-driven inflammation (11, 15, 16).

In contrast to exacerbating pathology in autoimmune conditions, Reg1-deficiency leads to improved resistance to at least one IL-17-mediated host response, disseminated candidiasis, causing enhanced survival and concomitantly reduced fungal kidney burdens (11). Therefore, although restraint of Reg1 is clearly beneficial to limit autoimmune pathology, it is conceivable that temporary blockade of Reg1 could be exploited to improve inflammatory immune responses during certain infectious settings. However, the impact of Reg1 in infectious disease settings has not been extensively explored.

KP is sensed by pattern recognition receptors such as TLR4, which activates a myeloid differentiation primary response gene 88 (MyD88)-dependent pathway leading to induction of pro-inflammatory chemokines and cytokines. TLR4 signaling also induces the TIR-domain-containing adaptor-inducing interferon-β (TRIF) pathway, which induces of Type I IFNs (IFN-α, IFN-β). Both MyD88 and TRIF signaling pathways are needed for defense against KP infection (17, 18). TLR4 is also upstream of IL-17 production from γδ T cells in response to KP (19), and *Il17ra*^−/−^ mice have uncontrolled KP infection with associated decreased expression of CXC chemokines, G-CSF and impaired neutrophil recruitment (20). Additionally, IL-17 synergizes with cytokines such as IL-22 to increase production of antimicrobial peptides in lung epithelium, such as LCN2, calprotectin (S100A8/9) and MUC1(21). Though produced by lymphocytes, IL-17 mediates downstream signals selectively on non-hematopoietic cells within the pulmonary epithelium during KP infection (22). Thus, the TLR4 and IL-17R pathways mediate an integrated inflammatory cascade that contains KP lung infection.

Given the documented role of Reg1 in restricting both TRL4- and IL-17-dependent signaling pathways, we hypothesized that Reg1-deficiency would be protective in the context of KP pneumonia, likely through decreased decay of pro-inflammatory mRNA transcripts. Indeed, we find here that Reg1-deficiency renders mice resistant to KP. However, bacterial burdens and typical pro-inflammatory cytokines known to be targeted by Reg1 were not altered in Reg1-deficent mice. Gene profiling of alveolar macrophages from KP-infected mice showed striking enrichment of type I IFN-associated gene pathways during Reg1 deficiency. Consistent with this, blockade of IFNR signaling reversed the protective effects of Reg1 deficiency. Mechanistically, Reg1 regulated the stability of mRNA encoding IRF7, an upstream regulator of Type 1 IFN gene expression by TLRs.

## Material and Methods

### Mice

*Reg1^+/−^* and *Reg1^+/+^ (Zc3h12a)* littermates on the C57BL/6 background were co-housed and used for all experiments. *Reg1(Zc3h12a)*^fl/fl^ mice are under material transfer agreement (MTA) from University of Central Florida (Orlando, Florida, USA). *Sftpc^Cre^* mice and CD45.1 mice were from The Jackson Laboratory. Mice were 8-12-weeks old and both sexes were used. All animal studies were approved by the Institutional Animal Care and Use Committee (IACUC) of the University of Pittsburgh.

### Bacterial infections and anti-IFNR1 treatment

KP ATCC strain 43816 was grown in tryptic soy broth to reach early log phase. 1-2×10^3^ CFU/mouse in PBS was administered by deep oropharyngeal aspiration. Where indicated, tamoxifen (TAM) was administered i.p. at 75 mg/kg dissolved in corn oil for 5 d and then rested for 5 d prior to induction of infection. *Reg1^+/−^* mice were treated with anti-IFNR1 antibodies or isotype control by i.p injection (Bio-Xcell MAR1-5A3, 250 ug/mouse, administered on Day 0 and Day 2 p.i). Mice were followed for 7 days.

### Bone marrow chimeras

WT (CD45.1) and *Reg1^+/−^* (CD45.2) mice were lethally irradiated (900 Gy). After 24h, each mouse received 4×10^6^ bone marrow cells by tail vein injection. Mice received antibiotics in drinking water starting 1 d before irradiation and continued for 10 d (trimethroprim/sulfametoxazole 200/40 mg). After 6 weeks, engraftment was confirmed by flow cytometry of blood for CD45.1 and CD45.2 markers.

### Bacterial burden, mRNA, and protein analysis

Tissues were homogenized in PBS and CFU levels were assessed by serial dilution plating. Lung tissues were homogenized in TrIzol (Invitrogen) and subjected to qPCR with SybrGreen probes. Threshold cycle (C_T_) values were normalized to *Gapdh.* ELISA kits were from eBioscience (Thermo Fisher Scientific) and R&D Systems. Abs used in western blots were cleaved caspase-3, total caspase 3 (Cell Signaling) and beta-actin (Abcam).

### Flow cytometry

Lungs were digested with lung dissociation kits (Miltenyi). Lung cell suspensions were stained for CD45, LyG, CD11b, CD11c, SiglecF, Ly6C, CD3, CD4, NK1.1, IFN-γ. Abs were from eBioscience, BD Biosciences or Biolegend. Alveolar macrophages (CD45^+^, LyG^−^ CD11b^−^ CD11c^hi^, SiglecF^+^) were isolated by FACS. Proliferation was assessed by BrDU incorporation (BD Biosciences); briefly, mice were injected i.p. with 1 mg of BrDU one day before sacrifice. Data were acquired on LSR Fortessa and analyzed with FlowJo software.

### Bone marrow derived macrophages (BMDM)

BM cells were extracted from *Reg1^+/−^* and *Reg1^+/+^* mice femur and tibia and differentiated into BMDM with L929 media (30%) for 5-6 days. Cultured BMDMs were stimulated with heat-killed KP (MOI = 10) up to 8 h or *Klebsiella* LPS (1 ug/ml) (Sigma-Aldrich).

### RNA Sequencing and Analysis

Samples from 12 individual mice (uninfected *Reg1^+/+^* n=3, uninfected *Reg1^+/−^* n=3, KP-infected *Reg1^+/+^* n=4, KP-infected *Reg1^+/−^* n=4) were used. Mice were infected with KP by oropharyngeal aspiration. After 24 h, alveolar macrophages were stained as described above and sorted by flow cytometry with a purity of 99%. Cells were placed into RLT-plus (Qiagen, Valencia, CA) and total RNA extracted using RNeasy MiniKits (Qiagen). RNA was quantitated using Nanodrop and integrity determined with a total RNA Nano Chip (Agilent Technologies). Single-stranded total RNA-seq libraries were sequenced with an Illumina Nextseq500 sequencer with a depth of 25 million reads per sample (75 bp single-end) at the University of Pittsburgh Health Sciences Sequencing Core. Fastq files with high quality reads (phred score >30) were uploaded to the CLC Genomics Workbench (Qiagen) and reads aligned to the mouse reference genome. Transcript counts and differential expression analyses were carried out using the CLC Genomics Workbench. RNAseq data were deposited to Sequence Read Archive (SRA), BioProject ID: PRJNA789160.

### Lung Histology

Lungs from *Reg1^+/−^* and *Reg1^+/+^* mice isolated at 48 h p.i. were fixed in 10% neutral buffered formalin for 24 h. Sections were stained with hematoxylin and eosin (H&E) by the University of Pittsburgh histology core. Slides were visualized on an EVOS microscope. Lung damage parameters measured were: A) Parenchymal inflammation and percentage of compromised tissue, B) peribronchial, and C) perivascular inflammation. The assigned score values were 0) none, 1) <25%, 2) 25-50%, 3) 50-75%, 4) >75%.

### Statistical Analysis

Data were analyzed on Prism (GraphPad). Data were analyzed by Log-Rank, One-way analysis of variance (ANOVA), Student’s t test, and post hoc tests were used as indicated. Each symbol represents one mouse unless indicated.

## Results

### Reg1-deficient mice are resistant to KP lung infection

A complete absence of Reg1 is extremely deleterious, as Reg1^−/−^ mice exhibit severe inflammation and severely shortened life spans (9). In contrast, *Reg1^+/−^* mice (sometimes termed here Reg1-deficient) have a normal life span without peripheral organ inflammation. Nonetheless, *Reg1^+/−^* mice show enhanced pro-inflammatory responses in multiple IL-17-driven autoinflammatory model settings (11, 15, 16). To determine whether Reg1 haploinsufficiency influences immunity to KP infection, *Reg1^+/−^* and *Reg1^+/+^* littermate controls were infected with KP by oropharyngeal aspiration. Survival and parameters of health including weight loss were monitored over time. Indeed, infection-induced survival and weight loss were significantly ameliorated in Reg1-deficient mice compared to *Reg1^+/+^* littermate controls (**Fig. 1A, B**). Unexpectedly, this improvement in survival was not accompanied by statistically significant differences in lung bacterial burdens (**Fig 1C**). Moreover, there were no changes in bacterial dissemination into the spleen between groups at any measured time point, even though *Reg1^+/−^* mice had lower expression of Reg1 (*Zc3h12a*) than controls **(Fig. 1D, E).**Furthermore, histologic analysis of infected lung tissues between *Reg1^+/+^* and *Reg1^+/−^* animals, showed that *Reg1^+/−^* mice developed smaller foci of pneumonia compared to controls, with particularly reduced levels of parenchymal inflammation (**Fig. 1F**, **G**). These data suggest that factors other than bacterial burden underlie the improved survival of Reg1-deficient animals.

**Figure 1:**
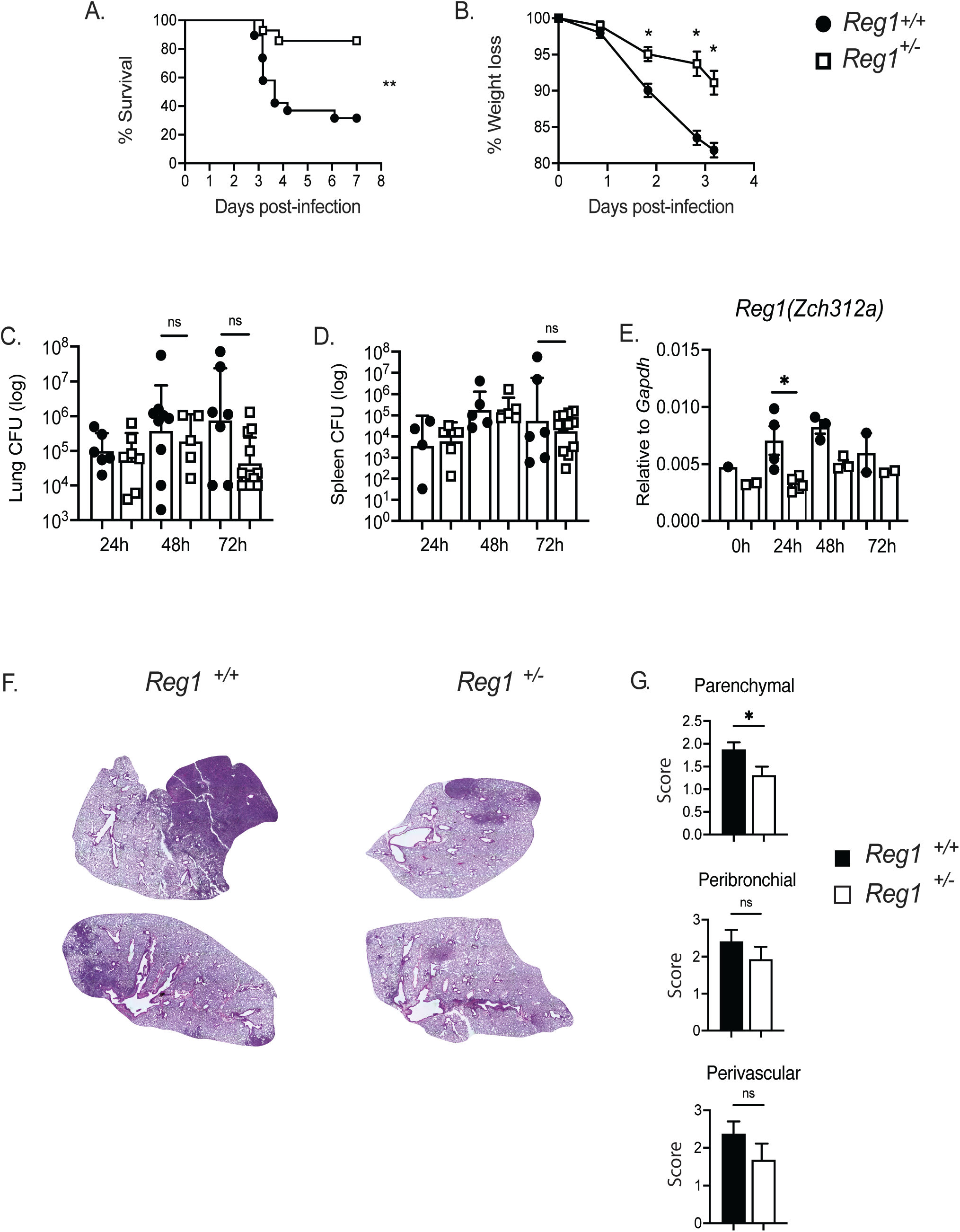
Reg1-haploinsufficient mice are resistant to KP lung infection. A) Lung and B) spleen bacterial burdens were assessed by CFU assessment at the indicated time points. Each symbol represents 1 mouse. Filled circle = *Reg1^+/+^* and open box= *Reg1^+/−^* mice. C) Weight loss was assessed at the indicated time points (n=15 mice/group). D) Survival of *Reg1^+/−^* or *Reg1^+/+^* littermates was assessed up to 8 days post-infection (n=15 mice/group). E) Reg1 gene expression (*Zc3h12a*) in lung homogenates of KP-infected mice. Data are pooled from two experiments. * P<0.05, **<0.01, Log-Rank test. F) Representative images of H&E lung sections from *Reg1^+/+^* and *Reg1^+/−^* mice at 48 h post-KP infection. G) Scoring of parenchymal, peribronchial and perivascular inflammation (*Reg1^+/+^ n=*6*, Reg1^+/−^* n=4). * P<0.05, unpaired-t test.

### Reg1 functions in both hematopoietic and non-hematopoietic compartments during KP pneumonia

Reg1 has been shown to act in multiple cell types, including T cells, macrophages and epithelial cells (7, 23). Of relevance to KP, Reg1 can restrict signaling by both TLR4 and IL-17. Whereas TLR4 acts predominantly on hematopoietic cells (especially macrophages), IL-17 predominantly mediates signaling in epithelial and/or mesenchymal target tissues, demonstrated in many settings including KP-infected lung (24). As a first step to define the essential compartments in which Reg1 functions to limit immunity to pulmonary KP infection, we used an adoptive transfer approach. Femoral bone marrow (BM) from *Reg1^+/−^* (CD45.2) or WT (CD45.1) mice were transferred into reciprocal irradiated *Reg1^+/−^* or WT recipients. After 8 weeks, engraftment was confirmed by flow cytometry (data not shown). Successfully reconstituted mice were infected with KP by oropharyngeal aspiration and followed up to 7 days. As expected, WT mice that received WT BM cells were susceptible to KP, whereas *Reg1^+/−^* mice receiving *Reg1^+/−^* BM were more resistant. As shown, *Reg1^+/−^* or WT mice that received *Reg1^+/−^* BM cells were resistant to KP, with almost 80% survival at day 7 post-infection. Additionally, *Reg1^+/−^* mice that received WT BM cells were resistant to KP, showing 60% survival at day 7 (**Fig 2A**). Thus, Reg1 appears to act in multiple compartments.

**Figure 2:**
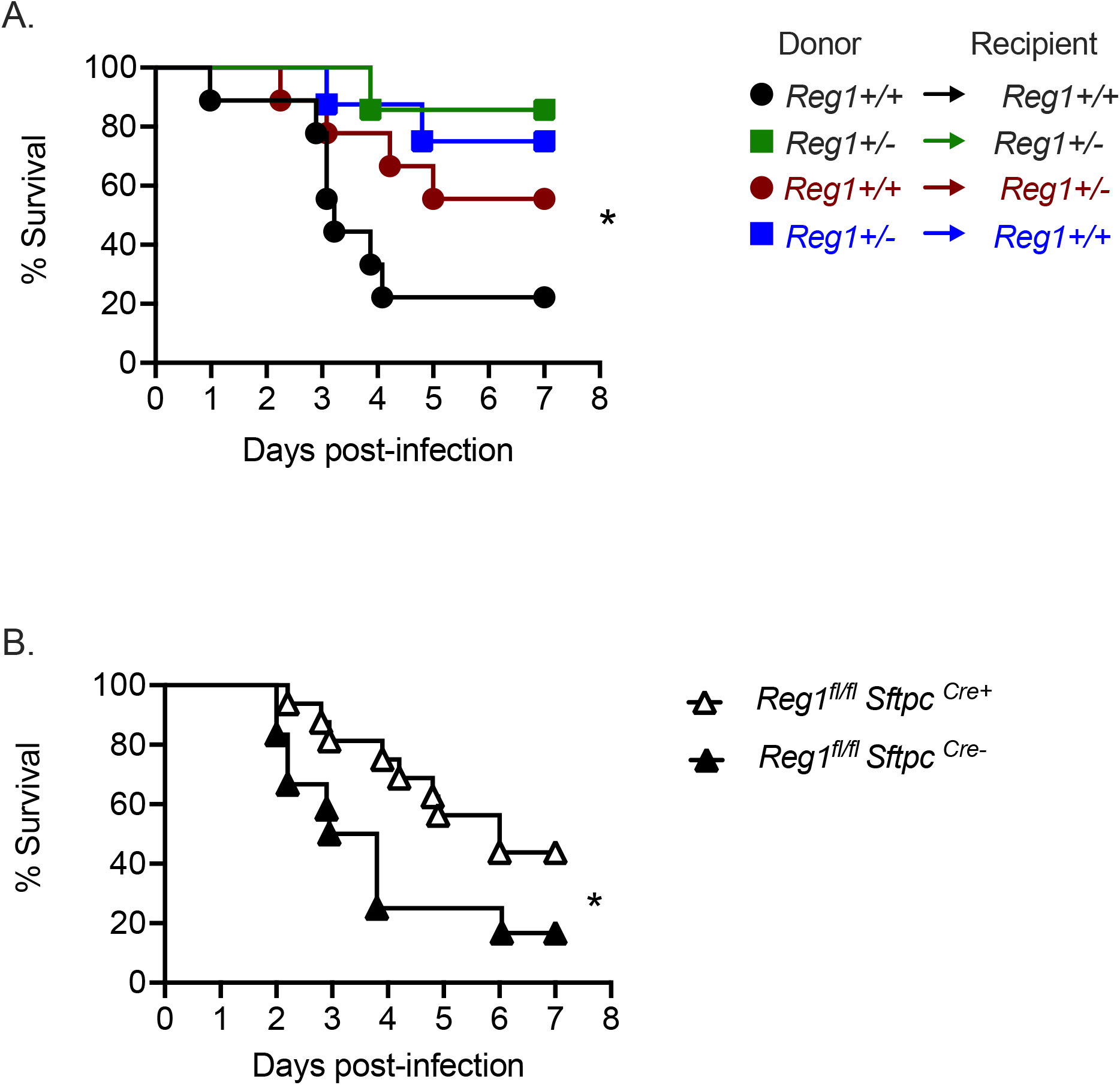
Reg1 deficiency in both hematopoietic and non-hematopoietic compartments contribute to prolonged survival upon KP lung infection. A) *Reg1^+/−^* and *Reg1^+/+^* littermates were irradiated to ablate bone marrow (BM) and later reconstituted with reciprocal femoral BM cells. Eight weeks after BM transfer, mice were infected with KP and survival monitored over 7 days. B) *Reg1^fl/fl^* were crossed to *Sftpc^CreERT2^* infected with KP and monitored over 7 d. Data are pooled from two experiments. * P<0.05, **<0.01, Log-Rank test.

Loss of IL-17RA in pulmonary epithelial cells renders mice highly susceptible to KP infection (24). Accordingly, given the potent impact of Reg1 on IL-17 signaling seen in prior studies (11, 12, 15, 25), we crossed *Reg1^fl/fl^* to surfactant protein C (*Sftpc)*^Cre^ mice in order to delete Reg1 conditionally in distal pulmonary epithelial cells (distal bronchi and alveoli). As shown, mice were modestly more resistant to KP than controls, though the improvement survival was much less profound than seen in *Reg1^+/−^* mice (**Fig 2B**), which is consistent with the BM chimera data showing contributions from both the hematopoietic and non-hematopoietic compartments contribute upon KP infection.

### Reg1 deficiency does not influence immune cell recruitment or proliferation during KP infection

Based on these data, it is evident that a Reg1 deficiency in the hematopoietic compartment provides a survival advantage to the host. We saw increased percentage of alveolar macrophages at 72h post-infection (**Fig. 3A**). However, there were no changes in the absolute numbers of recruited myeloid cells between *Reg1^+/−^* and control mice at 48 h and 72 h post-infection (**Fig 3B**). Consistent with this, expression of myeloid-recruiting chemokines such as *Cxcl1, Cxcl5* and *Ccl2* were similar among groups, despite the fact that these are all known transcriptional targets of Reg1 in other settings (**Fig. S1A**) (26, 27). There were also no differences in cellular proliferation and macrophage bacterial killing between *Reg1^+/−^* and control mice (**Fig 3B, C**). It is known that Reg1 controls B cell homeostasis and that complete Reg1 deletion in B cells increases antibody secretion (28). However, IgM and IgA levels in BALF and lung tissue were comparable between *Reg1^+/−^* and control mice (**Fig 3D**). Taken together, these data suggest that differences in survival during KP infection in *Reg1^+/−^* mice is not explained by altered recruitment of myeloid cells, cellular proliferation, macrophage killing capacity, or a B cell antibody response.

**Figure 3:**
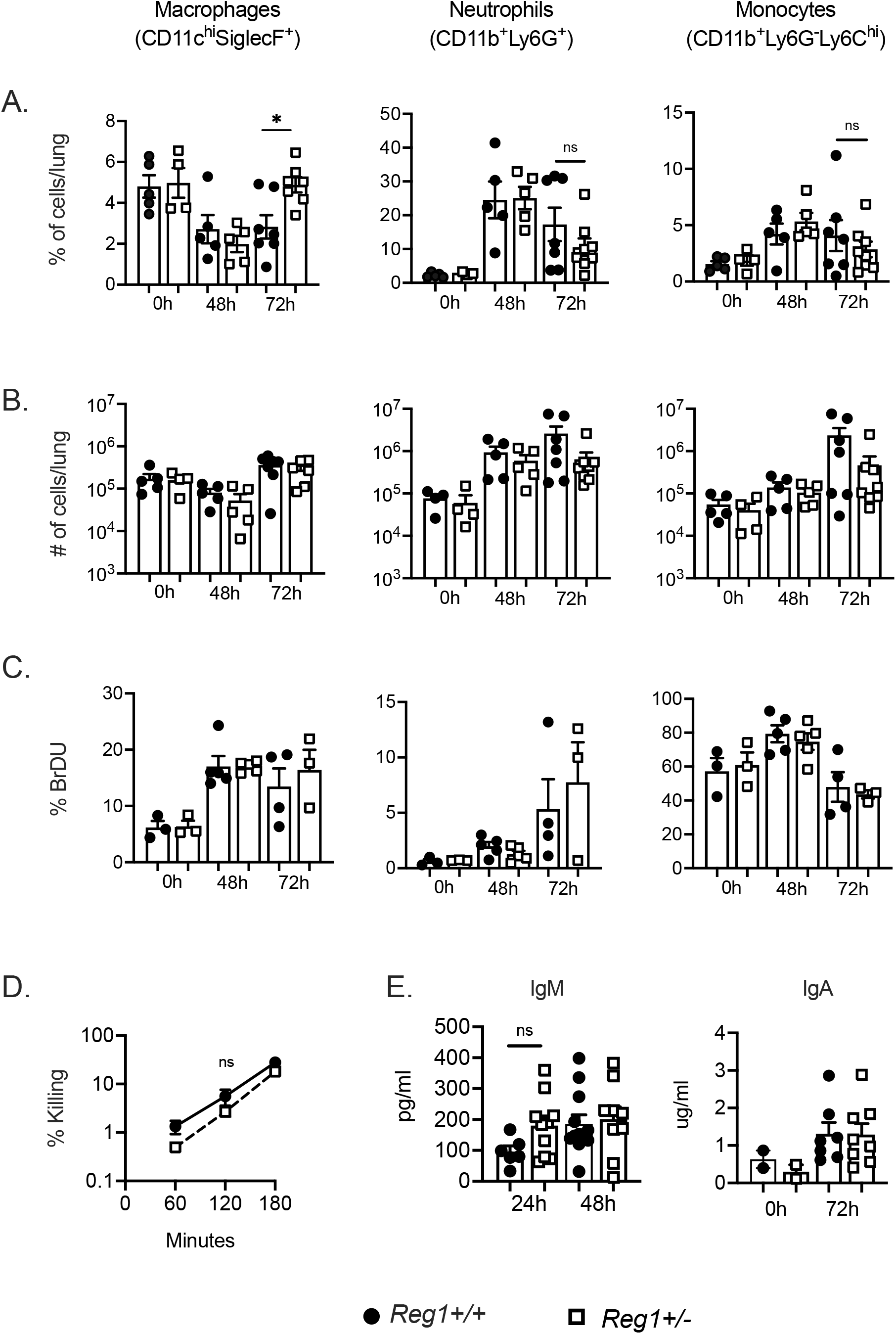
Immune cell recruitment, proliferation and antibody production are not affected by Reg1 deficiency during KP infection. A-B) Percentage and absolute numbers of cellular recruitment to the lung of infected mice up to 72h post-KP infection. C) BrDU incorporation of lung myeloid cell populations were assessed by flow cytometry at baseline (0 h) or 48 and 72 h post-infection. Data are pooled from two independent experiments (A-C). D) Macrophage killing and Ab production in lung was assessed at the indicated time points. Data are pooled from two independent experiments. Data analyzed by one-way ANOVA *P<0.05, ns: not significant.

### Resistance to KP caused by Reg1 deficiency is linked to increased interferon (IFN) signature

To understand the mechanisms by which Reg1 deficiency promotes survival, we performed transcriptomic profiling of purified alveolar macrophages from *Reg1^+/−^* and littermate control lungs at 24 h post-KP infection. This time point was chosen to define the early events that are operative during the initial stages of infection. There were 796 differentially expressed genes, of which 237 were upregulated and 559 were downregulated. Gene set enrichment analysis (GSEA) revealed major changes in type I IFN-related responses (IFN-α), oxidative phosphorylation, and IFN-γ response as the top three enriched gene sets, with a normalized enrichment score (NES) <2.5 for the type I IFN group (**Fig. 4A, B**).

**Figure 4:**
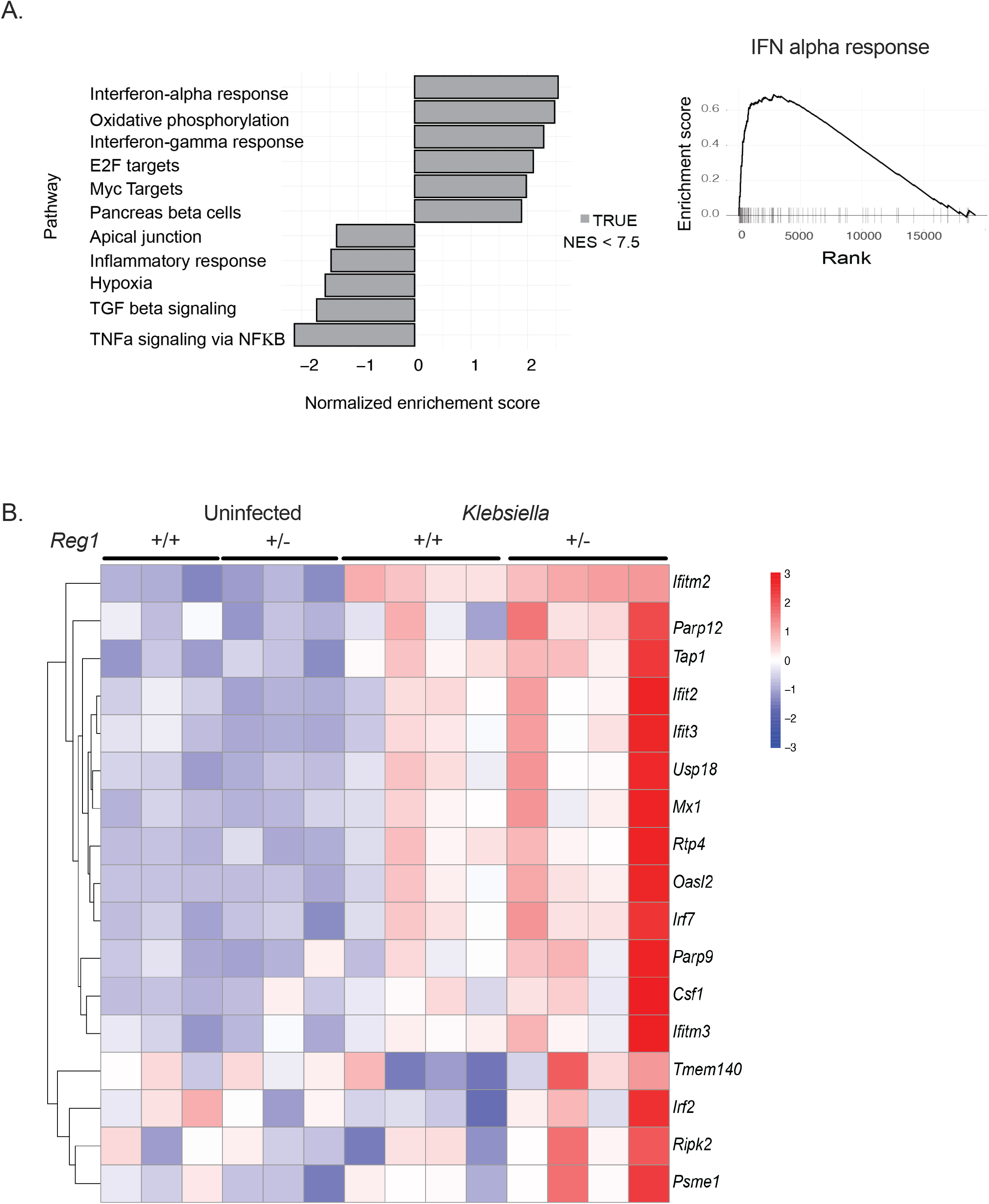
RNASeq of sorted alveolar macrophages reveals an enriched Type 1 IFN gene signature. *Reg1^+/−^* and *Reg1^+/+^* littermates were infected with KP or PBS by oropharyngeal aspiration. Alveolar macrophages were sorted by flow cytometry at 24 h post infection and subjected to RNASeq. A) Gene set enrichment analysis (GSEA) of type I IFN gene sets. B) Heatmap depicting differentially expressed genes in the type I IFN pathway (n=4 mice/group).

We next validated these pathways by functional analysis. Consistent with the RNASeq data, Type I IFN (IFN-β) expression was higher in lung homogenates of Reg1-deficient mice compared to controls at 24 h and 48 h post-infection **(Fig 5A**). Since IFN-β can be produced by macrophages during Gram-negative pneumonia (29), we stimulated *Reg1^+/−^* or control BMDMs with heat-killed KP and assessed *Ifnb1* mRNA. Indeed, *Ifnb1* was elevated in Reg1-deficient macrophages compared to controls as early as 2 h post-infection (**Fig. 5B**). In contrast, IFN-γ expression was similarly expressed between *Reg1^+/−^* and Reg1^+/+^ mice after KP infection (**Fig 5C**). Similarly, oxidative phosphorylation, of sorted alveolar macrophages assessed by mitochondrial respiration, was not different between *Reg1^+/−^* mice and controls (**Fig. S1B,C**). Collectively, these data indicate that Reg1 deficiency induces IFN-β upon KP infection.

**Figure 5:**
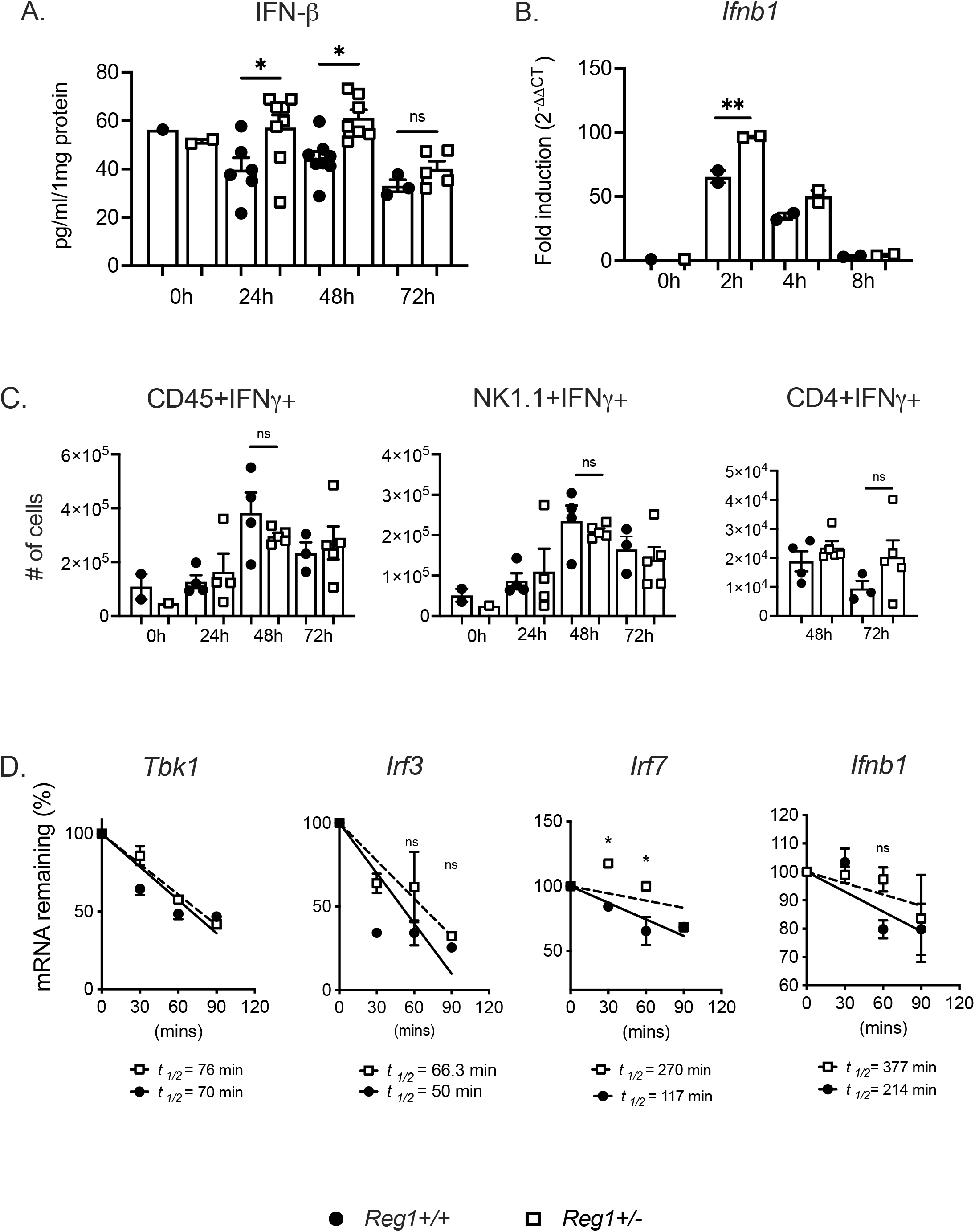
Increased IFN-β but not IFN-γ in Reg1 deficient mice during KP infection. *Reg1^+/−^* and *Reg1^+/+^* littermates were infected with KP. A) IFN-#x03B2; was assessed in lung homogenates by ELISA. B) *Ifnb1* mRNA was assessed in cultured BMDMs infected with heat-killed KP for the indicated time points. C) Levels of IFN-γ were assessed by intracellular staining in total CD45^+^, NK1.1^+^ and CD4^+^ cells. B) Each symbol represents one mouse. *P<0.05 One-way ANOVA with post-hoc Tukey’s test. D) BMDMs were pretreated with LPS for 3 h and treated with Act D to stop new transcription. Expression of the indicated mRNAs were assessed for the indicated times by qPCR, normalized to *Gapdh*. Levels compared to time = 0 data are presented as means ± SEM, representative of 2 independent experiments. One-way ANOVA with post hoc Tukey’s test *P < 0.05 by; RNA half-life (*t*_½_) was determined by linear regression as described (50).

Reg1 functions primarily by promoting endonucleolytic decay of target mRNA transcripts (30). Based on the observed increased *Ifnb1* expression in Reg-deficient BMDMs, we hypothesized that Reg1 deficiency may result in prolonged stabilization of *Ifnb1* mRNA or regulatory factors upstream of *Ifnb1* regulation. To test this, *Reg1^+/−^* and control BMDMs were stimulated with LPS to prime expression of genes in the IFN pathway. Cells were treated with actinomycin D (Act D) to block further transcription, and the half-life of candidate target transcripts was assessed over a 2-hour time course. Although the intrinsic mRNA stability of *Ifnb1* or other mRNAs was not detectably altered in Reg1-deficient cells, the half-life of *Irf7* mRNA considerably increased in the setting of Reg1 deficiency (**Fig. 5D**), potentially explaining the increased IFN-β seen in Reg-1 deficient mice upon KP pneumonia.

To determine whether Type I IFN accounted for the prolonged survival in *Reg1^+/−^* mice, we administered anti-IFNR1 Abs or isotype controls at Day 0 and 2 post-KP infection and followed mice for 7 days. Strikingly, 100% of *Reg1^+/−^* mice that received isotype control Abs survived, while 60% of *Reg1^+/−^* mice that received anti-IFNR1 succumbed (**Fig. 6**). Therefore, the protection afforded by Reg1-deficiency requires Type I IFN signaling.

**Figure 6:**
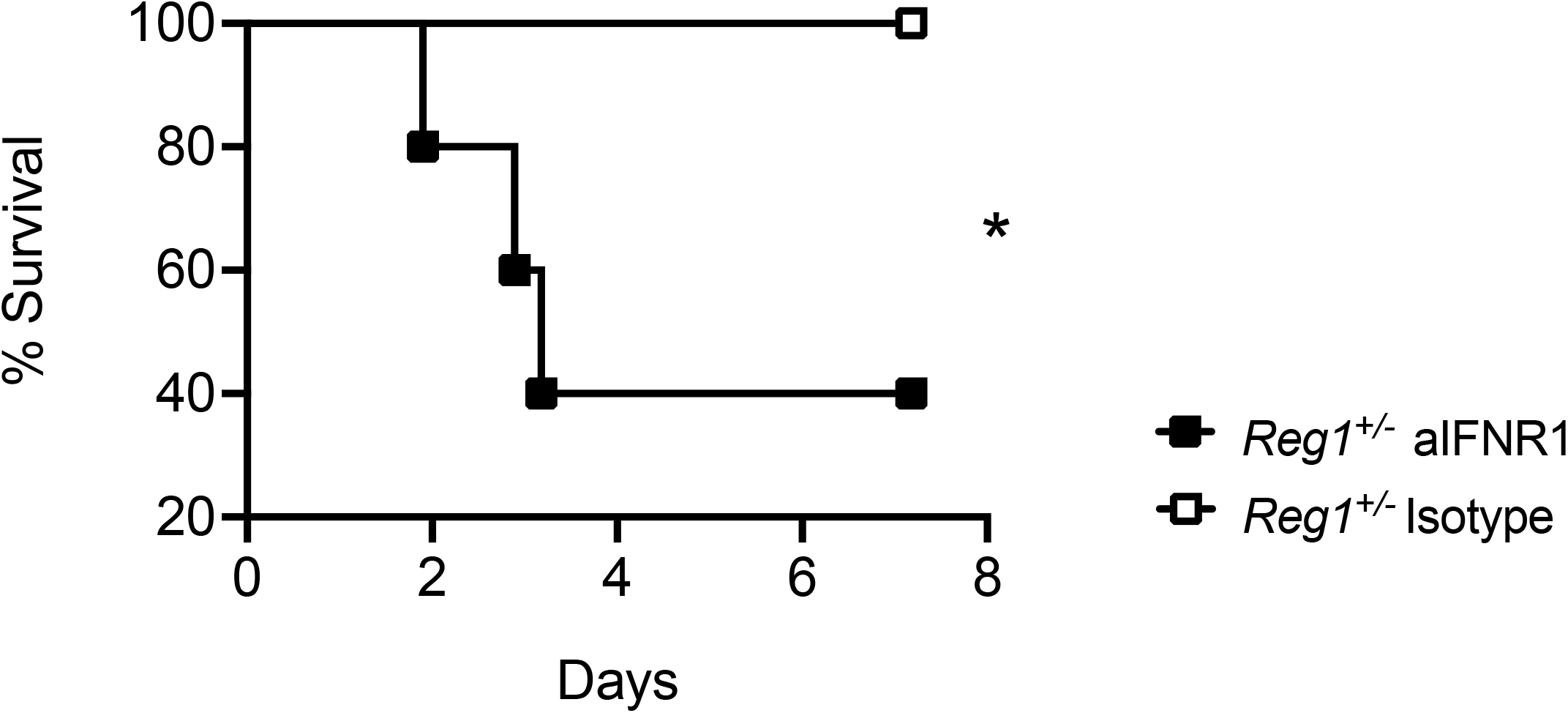
IFNR1 blockade in Reg1-deficient mice enhances susceptibility to KP infection. *Reg1^+/−^* mice received anti-IFNR1 antibody or isotype control by i.p injection on Day 0 and Day 2 post-KP infection. Mice were followed up to 7 days post-infection. (n=5 mice/group). * P<0.05, Log-Rank test.

### Increased Ifnb1 correlates with enhanced anti-inflammatory response in Reg1-deficient mice

IFN-β has the capacity to exert anti-inflammatory effects in the host, reported in several disease models. This effect has been attributed in part to type I IFN induction of IL-10, which dampens expression of transcription of cytokines such as *Tnfa, Cxcl1, Il6* (31, 32). Based on this, we assessed IL-10 in lung homogenates of KP-infected mice. There was a trend to increased IL-10 at 72 h post-infection in *Reg1^+/−^* mice compared to control mice, though this did not reach statistical significance (**Fig 7A**). TNF-α was not affected *Reg1^+/−^* compared to controls (**Fig 7B**). We then asked whether apoptosis was affected in *Reg1^+/−^* mice compared to controls. As shown, cleaved caspase 3 expression in total lung homogenates was increased in *Reg1^+/−^* mice compared to controls (**Fig 7C, D**). These data suggest that increased IFN-β arising from Reg1 deficiency may confer an anti-inflammatory advantage to *Reg1^+/−^* mice sufficient to prolong survival upon KP infection.

**Figure 7:**
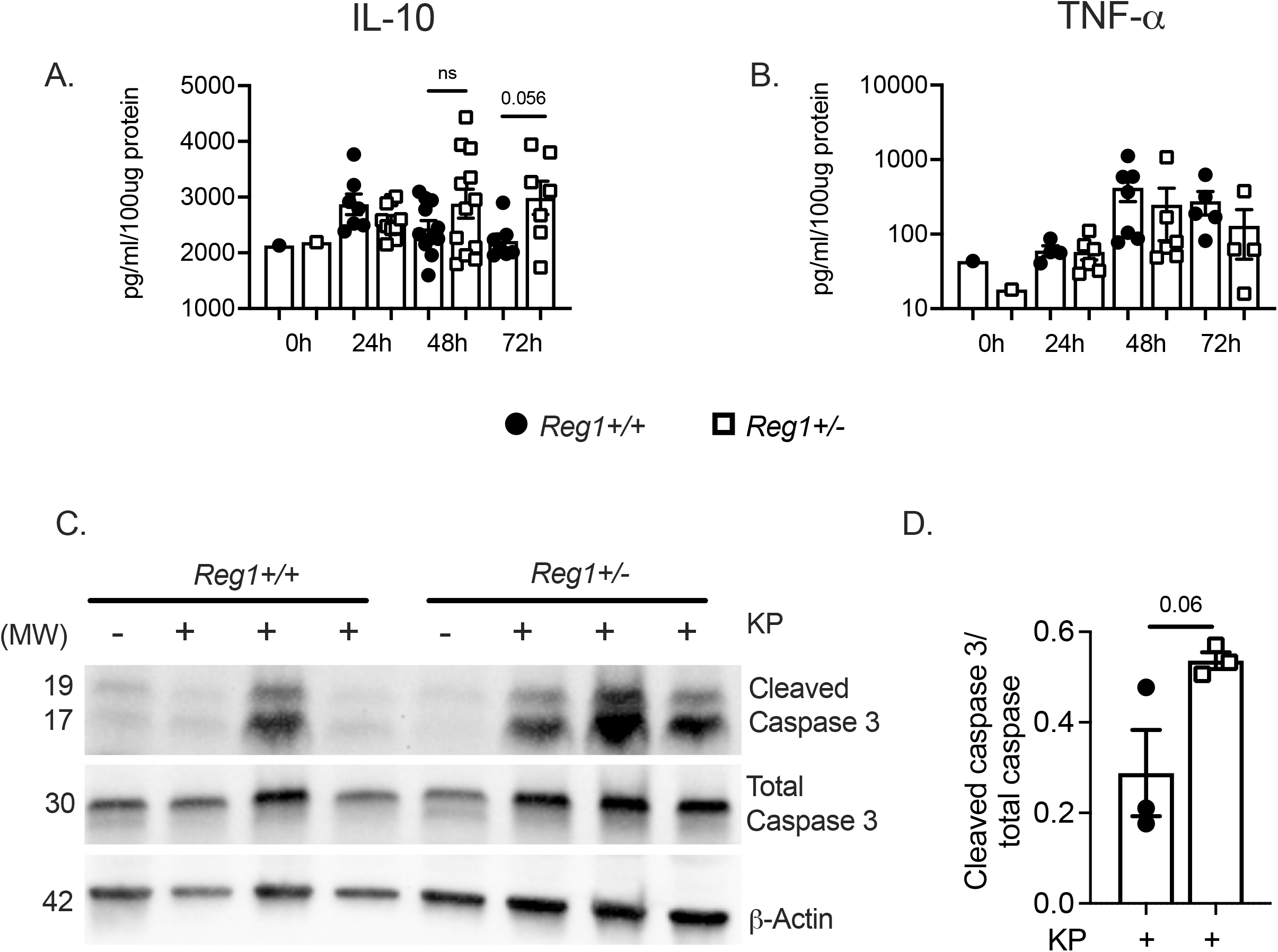
Reg1 deficiency enhances the inflammatory response and increases apoptosis. A, B) IL-10 and TFNα were assessed by ELISA in lung homogenates of the indicated mice infected with KP. Data are pooled from two independent experiments. C) *Reg1^+/−^* and *Reg1^+/+^* littermates were infected with KP. Mice were sacrificed at 48h post-infection, and cleaved Caspase 3 and total Caspase were assessed in lung homogenates by immunoblotting. At right, quantified band intensity values ± SEM is shown. Data are representative of individual replicates.

## Discussion

Reg1 controls the inflammatory response by degrading many individual inflammatory cytokine mRNAs, including *Il6, Il1b,* and *Il12b,* thus tempering the overall inflammatory milieu (9). Perhaps even more significantly, Reg1 degrades transcripts encoding inflammatory transcription factors (TF) such as *Nfkbiz,* which encodes the non-canonical TF IκBξ, and therefore Reg1 can indirectly affect all IκBξ-inducible genes (11). Thus, it is not unexpected that a complete knockout of Reg1 gene (*Zc3h12a*) is fatal due to severe autoinflammation. *Reg1^+/−^* mice, on the other hand, appear to have a normal life span, fertility, and do not exhibit peripheral baseline inflammation. However, *Reg1^+/−^* mice still have an enhanced inflammatory response, particularly when the disease model relies heavily on the IL-17R pathway (11, 15, 25). In this KP infection model, we were surprised that levels of inflammatory cytokines and chemokines were similar between Reg1-deficient mice and controls. Since these mice are haploinsufficient for the Reg1 gene (*Zc3h12a*), and KP is sensed by TLR4, one potential explanation is the remaining Reg1 levels may be enough to compensate for the cytoplasmic MyD88-driven pathway, but not the endosomal TRIF-dependent pathway, thus leading to increased type I IFN expression and protection from KP.

Since most pro-inflammatory genes known to be driven by Reg1 were unchanged in lung tissues regardless of Reg1 deficiency, and there was a clear increase in the percentage of alveolar macrophage (AM) recruitment, we performed RNAseq in AMs cells in order to evaluate macrophage-intrinsic activities underlying the Reg1-deficient phenotype. This approach has the caveat that contributions from other cells such as interstitial macrophages, inflammatory monocytes or other immune cells may be missed. In this regard, CCR2+ inflammatory monocytes recruited during KP infection are known to enhance IL-17 production from ILC3 cells leading to KP eradication (33, 34). However, this mechanism has been shown to be important with clinical strains of KP and not with the serotype used here (ATCC 43816). Future studies will focus on additional cell types to determine more broadly where Reg1 is operative in this setting.

The role of type I interferons (IFN-α/IFN-β) in bacterial infections is not fully defined, especially when compared to its extensively-studied roles in viral infections. Even among bacterial pathogens, the response to type I IFN differs depending on site of infection and pathogen characteristics. For instance, type I IFN is detrimental in animal models of *Staphylococcus aureus* and *Listeria monocytogenes* (35, 36). On the other hand, there is a protective response in models of *Legionella pneumophilia* and *Streptococcus pyogenes (37).* In the case of *Klebsiella pneumoniae,* IFN-β signaling in NK cells enhances secretion of IFN-γ, decreasing bacterial burden (38). In this study, we did not see differences in recruitment of NK cells or enhanced IFN-γ secretion by NK cells in *Reg1^+/−^* mice compared to controls. However, the enhanced mortality in control WT mice is significantly decreased in *Reg1^+/−^* mice, which we propose is due to increased type I IFN that modulates the inflammatory response enough to facilitate bacterial eradication and pneumonia resolution, and this model is supported by histological evidence as well as the observation that anti-IFNR1 blockade reverses the protection provided by Reg1 deficiency.

IFN-β can lead to many complex events in bacterial infections, including increased apoptosis, macrophage efferocytosis and resolution of infection in models of *E. coli* pneumonia and peritonitis, associated with enhanced IL-10 secretion (29). IL-10 mediates many anti-inflammatory effects and regulates metabolic reprograming of macrophages (39). Related to this, interstitial macrophages have immunoregulatory properties by secreting IL-10 upon LPS and CPG-DNA stimulation in models of asthma (40). Although there was only a modest trend of increased IL-10 in these settings, there was a clear increase in cleaved Caspase-3 in Reg1-deficient lungs, correlating with increased IFN-β and resistance to KP pneumonia. Concomitantly, transcriptomic profiles revealed several IFN-stimulated genes (ISGs) implicated in regulating apoptosis, including *Ifit2, Ifit3* and *Ifitm3*. Intriguingly, IFIT2 is an RBP that enhances apoptosis (41, 42), though its precise role in the context of Reg1 immunoregulation is as yet unknown. Thus, we speculate one mechanism underlying these results is through actions of type I IFN mediating increases in apoptosis, an important step for resolution of pneumonia (43).

Since IFN-β levels were increased in Reg1-deficient mice upon KP infection, we initially expected that Reg1 deficiency would result in enhanced stability of the *Ifnb1* transcript. However, our data instead indicate that the impact on *Ifnb1* appears to be indirect via control of its upstream regulator *Irf7* mRNA, opening a new facet of how Reg1 influences immune responses. A deficiency in a related RBP, Regnase-3 (*Zc3h12c*) also leads to increased IFN (type I and II) signaling, and IRF7 can transcriptionally control expression of Regnase-3 (44). However, Reg3 expression was not altered in AMs based on the RNAseq data or in total lung by qPCR, and thus we believe this axis is not a major driver of the effects seen here (**Fig. S2D**).

The Type III interferons have emerged in recent years as key regulators of antiviral and antifungal immunity (45). It has been shown that IFN-lambda (L) increases lung epithelial permeability and facilitates bacterial transmigration in animal models infected with KP (KP ST258), an effect that is counteracted by IL-22 (46). In this model we did not see increases in IFN-L despite increase of type I IFN and in vitro stability of IRF-7, though determining whether this axis of interferon activity contributes in any way will require further study (**Fig. S2C**). Surprisingly, though IL-22 and IL-17 signaling are well described to play important roles in KP eradication (20, 21, 24), their expression was also not altered by Reg1 deficiency (**Fig. S2A,B**). This would be consistent with activities of Reg1 being downstream of these or other cytokines rather than upstream of their production, though further analyses will be required to prove that point definitively.

The immune system has evolved to balance the vital effects of anti-microbial effector functions with the potential for causing collateral tissue damage. In this regard, the activation of every immune signaling pathway is accompanied by negative feedback signaling events that restrain inflammation (47, 48). In selected conditions, however, it may be clinically beneficial to allow more fulminant inflammation in order to treat a life-threatening condition. Indeed, checkpoint inhibitor blockade for cancer therapy is built on exploiting this concept (49). By analogy, releasing inflammatory “brakes” may be useful in the context of severe infections such as bacterial pneumonia, and the present data suggest that Reg1 could, in principle at least, be one such target.

## Acknowledgements

GTN was supported by NIH grants HL135476, HL154231, AI153549. KC was supported by HL137709. SLG was supported by AI147383. We thank BM Coleman for technical support and JK Kolls for helpful discussions. We thank the reviewers for excellent suggestions that significantly improved the mansucript.

## Abbreviations

AM: alveolar macrophage
BM: bone marrow
BMDM: bone marrow-derived macrophages
KP: *Klebsiella pneumoniae*
MCPIP1: MCP1-induced protein 1 (alternative name for Regnase-1)
MyD88: myeloid differentiation primary response gene 88
RBP: RNA binding protein
Reg1: regnase-1
TAM: tamoxifen
TLR: Toll-like receptor
UTR: untranslated region
VAP: ventilator-associated pneumonia

## Author contributions

Conceptualization – GTN, PSB, SLG, KC

Investigation – GTN, BL, LF, FEYA, KC

Writing – GTN, SLG

Reviewing and Editing – GTN, KC, PSB, SLG

Visualization – GTN

Supervision – GTN, SLG, PSB

Funding Acquisition – GTN, SLG, KC, PSB

---

**Figure S1: Expression of chemokines and mitochondrial respiration is similar between Reg1^+/−^ and controls**. *Reg 1^+/−^* and *Reg1^+/+^* littermates were given PB or KP by oropharyngeal aspiration and sacrificed at 24h. A) *Cxcl1, Cxcl5* and *Ccl2* expression was measured in lung tissue. Data analyzed by Student t-test and found non-significant B) CD11c^+^ cells sorted from PBS-treated or C) KP-infected mice were sorted and evaluated for oxygen consumption rate (OCR) by Seahorse flux analyzer (n=4 mice per group). Data analyzed by ANOVA and found to be non-significant.

**Figure S2: Expression of IL-17, IL-22 and IFN-λ are similar between *Reg1^+/−^* and controls.** *Reg 1^+/−^* and *Reg1^+/+^* littermates were infected with KP and sacrificed at the indicated time points. Lung tissues were assessed for protein and mRNA. A) IL-17, B) IL-22, C) IFN-λ, expression in lung homogenates, D) *Regnase-3 (Zc3h12c)* mRNA expression in total lung. Data are pooled from two independent experiments. Data analyzed by ANOVA and found to be non-significant.

## Notes

### Competing Interest Statement

SLG has consulted for Eli Lilly and Aclaris Therapeutics in the past 3 years

### Summary of Updates

Several new experiments have been added to address the role of Type I interferons in Regnase-1 mediated protection from Klebsiella pneumoniae infections in mice

