## Supplementary figures and images for "Regnase-1 deficiency restrains *Klebsiella pneumoniae* infection by regulation of a Type I interferon response"

### Supplemental Figs 1-2

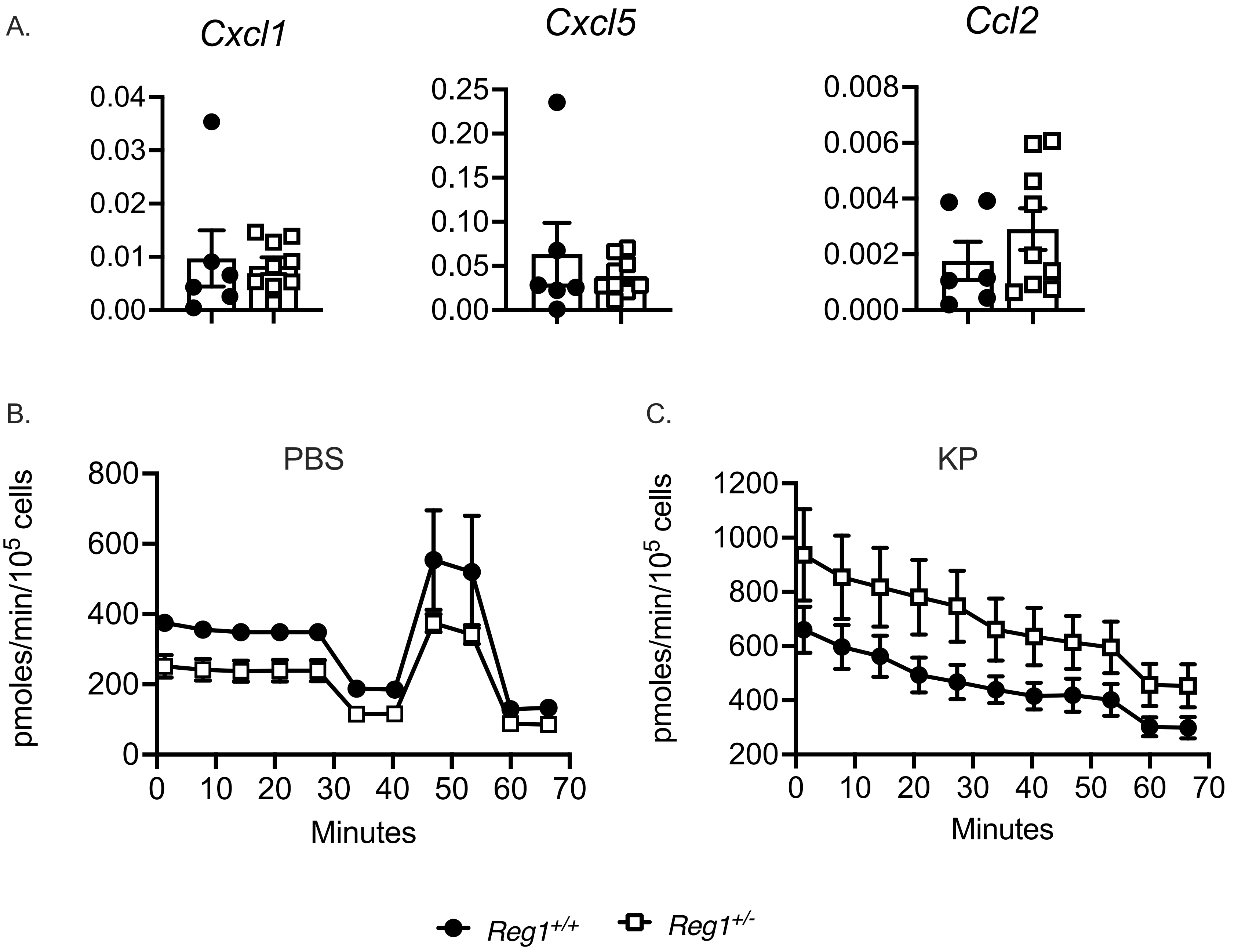

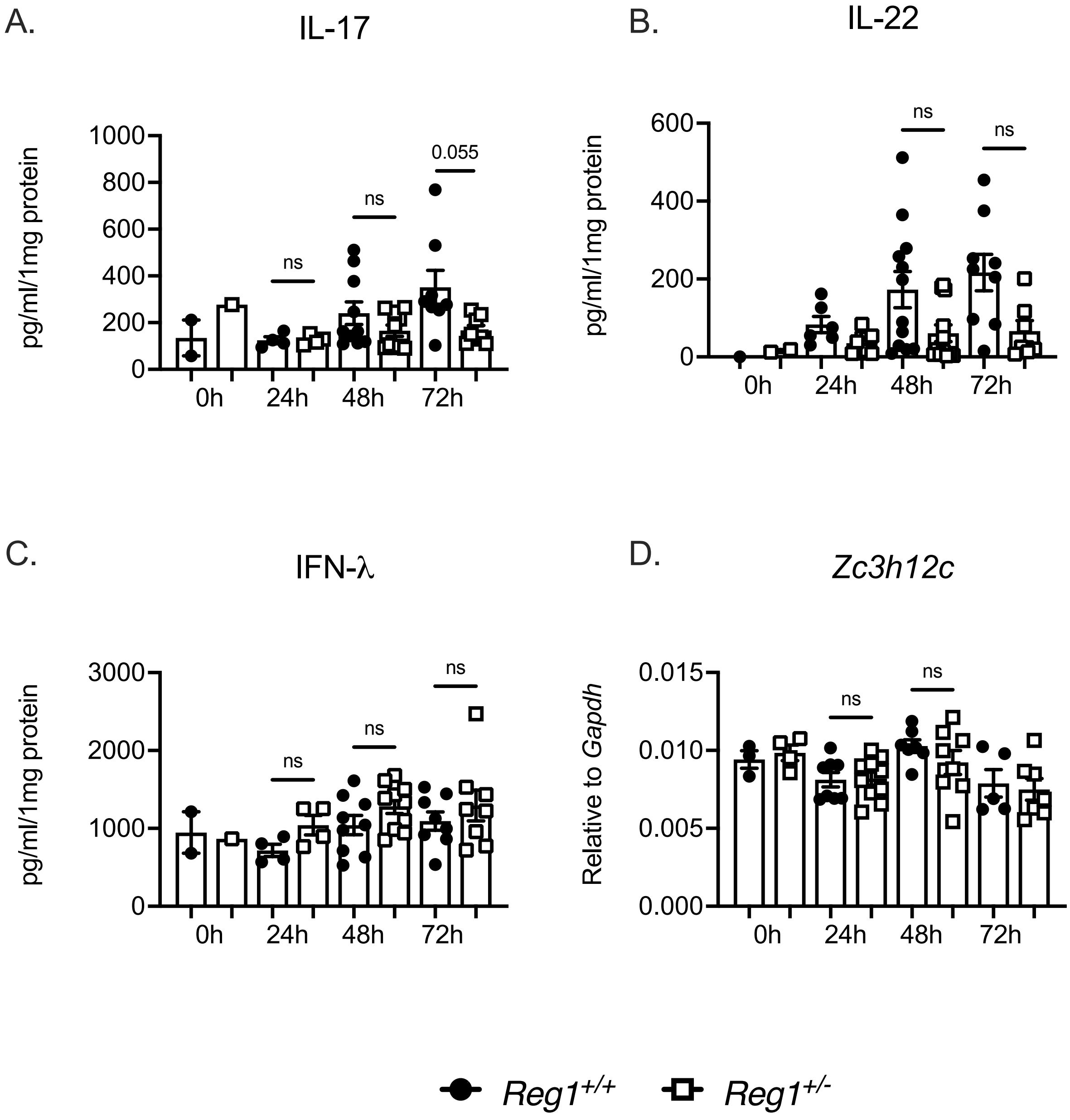
